## Supplementary figures and images for "Lipidome visualisation, comparison, and analysis in a vector space"

### S1 Fig

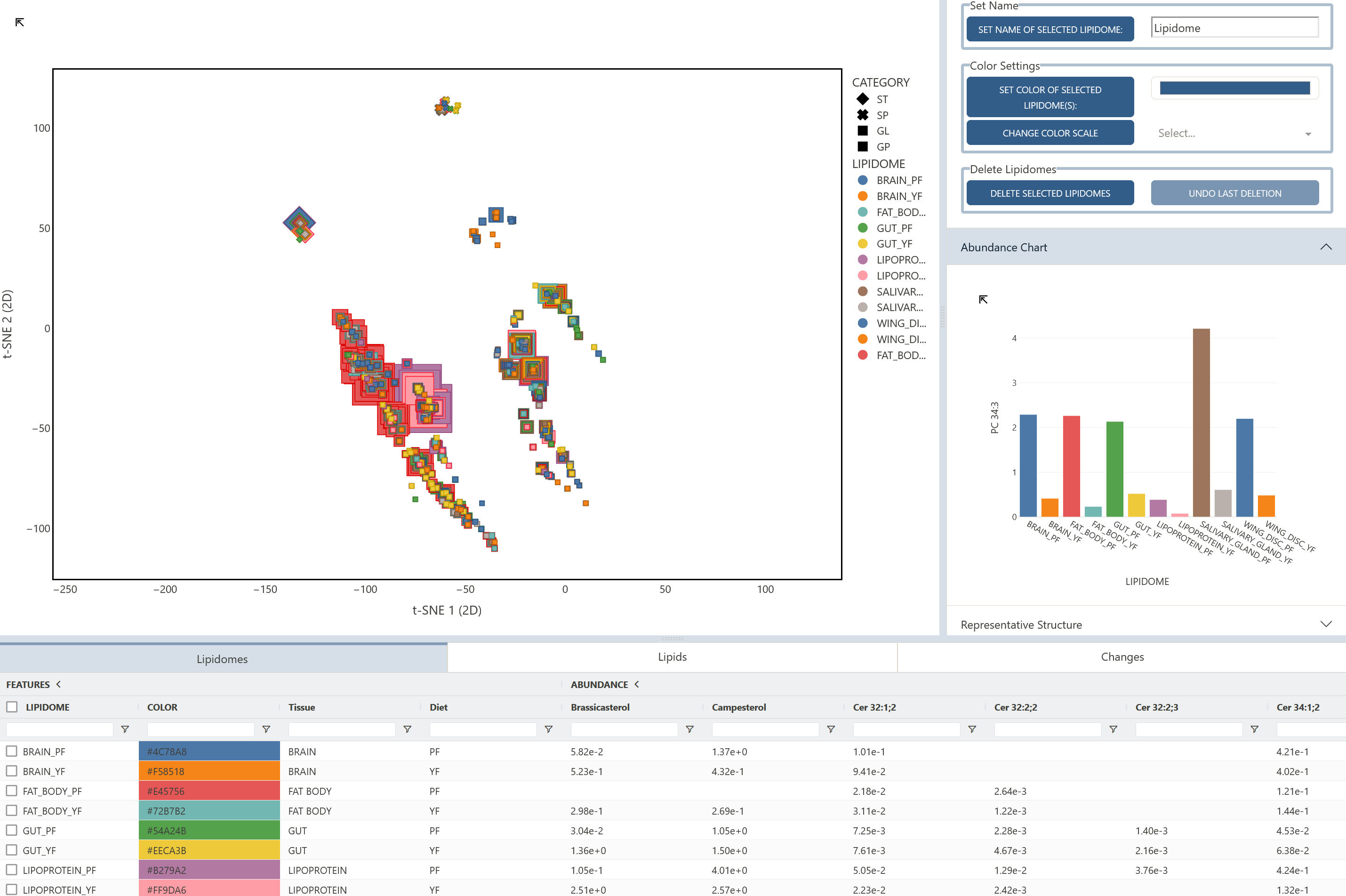

### S2 Fig

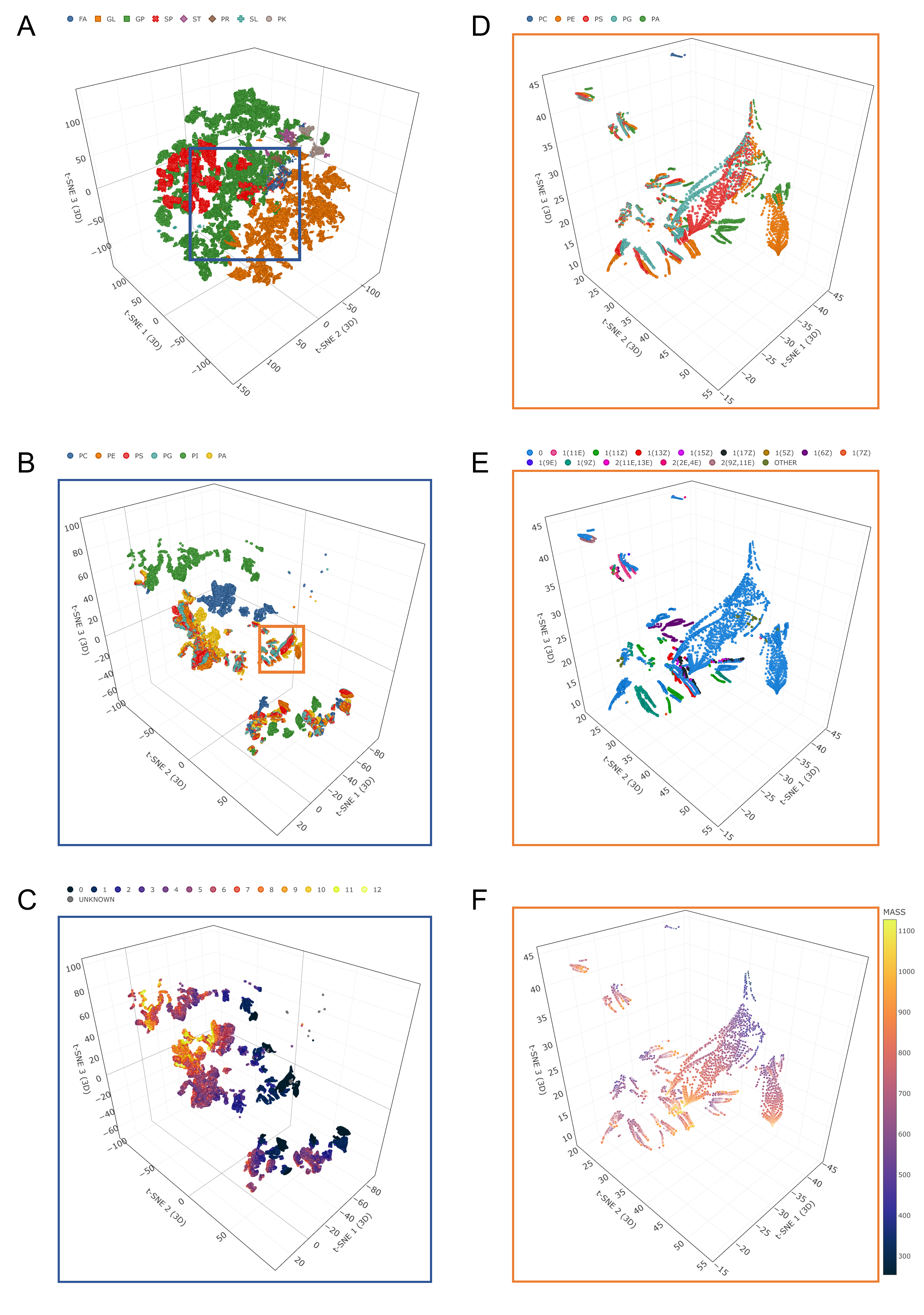
