## Supplementary material for "Lipidome visualisation, comparison, and analysis in a vector space": S1 Table

**S1 Table. Word2Vec Embedding Parameters.**

| Word2Vec | | | | | |
| --- | --- | --- | --- | --- | --- |
| Vector size | Context window | Architecture | Optimization | Minimum word frequency | Training iterations |
| 100 | 4 | Skip-gram | Hierarchical softmax | 1 | 10 |
