## Supplementary material for "Lipidome visualisation, comparison, and analysis in a vector space": S2 Dataset: drosophila_dataset_derivation.py

```
from pathlib import Path

import pandas as pd

drosophila_original_df: pd.DataFrame = pd.read_excel(
    "msb201229-sup-0002.xls",
    sheet_name="2.Larval tissues",
    header=None,
)

drosophila_transformed_df: pd.DataFrame = drosophila_original_df.iloc[
    3:, 1:
].reset_index(drop=True)

drosophila_transformed_df.iloc[0:2, :] = drosophila_transformed_df.iloc[
    0:2, :
].fillna(
    method="ffill", axis="columns"
)

drosophila_transformed_df = drosophila_transformed_df.loc[
    :,
    (
        (drosophila_transformed_df.iloc[1] != "Standard Deviation")
        & (
            ~drosophila_transformed_df.iloc[2]
            .str.contains("LDF")
            .fillna(False)
        )
    ),
]

drosophila_transformed_df.replace("Stigmsterol", "Stigmasterol", inplace=True)

tissue_dfs: list[pd.DataFrame] = []

for group in drosophila_transformed_df.T.groupby(0):
    tissue_df: pd.DataFrame = group[1].T
    columns: pd.Series = tissue_df.iloc[0:3].apply(
        lambda column: (
            column.iloc[1]
            if column.iloc[1] == "Lipid species"
            else column.iloc[0].replace(" ", "_") + "_" + column.iloc[2]
        )
    )
    tissue_df.columns = columns
    tissue_df = (
        tissue_df.iloc[3:]
        .dropna(how="all")
        .reset_index(drop=True)
        .set_index("Lipid species")
    )

    tissue_dfs.append(tissue_df)

drosophila_transformed_df = pd.concat(tissue_dfs, axis="columns").T

drosophila_transformed_df.index.name = "LIPIDOME"

drosophila_transformed_df = drosophila_transformed_df.sort_index(
    axis="index"
).sort_index(axis="columns")


def adjust_drosophila_lipid_name(name: str) -> str:
    if "Cer" in name:
        components: list[str] = name.split(":")
        name = ":".join(components[0:2]) + ";" + components[2]

    return name


drosophila_adjusted_lipid_names: pd.Series = (
    drosophila_transformed_df.columns.to_series().apply(
        adjust_drosophila_lipid_name
    )
)

drosophila_transformed_df.columns = drosophila_adjusted_lipid_names

drosophila_abundances_df: pd.DataFrame = drosophila_transformed_df

drosophila_features_df: pd.DataFrame = (
    drosophila_transformed_df.index.to_series()
    .str.rsplit("_", expand=True, n=1)
    .rename(columns={0: "Tissue", 1: "Diet"})
)

drosophila_features_df["Tissue"] = drosophila_features_df["Tissue"].str.replace("_", " ")

output_dir: str = "../lipidome_projector_datasets"

Path.mkdir(Path(output_dir), exist_ok=True)

drosophila_abundances_df.to_csv(output_dir + "/drosophila_abundances.csv")
drosophila_features_df.to_csv(output_dir + "/drosophila_features.csv")
```
