## Supplementary material for "Lipidome visualisation, comparison, and analysis in a vector space": S2 Dataset: lamp3_dataset_derivation.py

```
from pathlib import Path

import pandas as pd

lamp3_original_df: pd.DataFrame = pd.read_excel(
    "journal.pgen.1009619.s007.xlsx",
    sheet_name="BAL mol%",
)

lamp3_transformed_df: pd.DataFrame = (
    (
        lamp3_original_df.drop(columns=lamp3_original_df.columns[1:13])
        .set_index("LipidSpecies")
        .T.rename_axis("LIPIDOME", axis="index")
    )
    .sort_index(axis="index")
    .sort_index(axis="columns")
)

lamp3_transformed_df = lamp3_transformed_df.rename(
    columns={"FC": "ST 27:1;1"}
).sort_index(axis="columns")


def adjust_sp_names(name: str) -> str:
    if name.startswith("Cer") or name.startswith("SM"):
        return name.replace(";0", "")
    else:
        return name


lamp3_transformed_df.columns = pd.Index(lamp3_transformed_df.columns.to_series().apply(adjust_sp_names))


def determine_lamp3_features(name: str) -> dict[str, str]:
    feature_code: str = name.split("_")[1]
    return {
        "Genetics": "WILDTYPE" if feature_code.endswith("WT") else "LAMP3-KO",
        "Challenge": "ASTHMA" if feature_code.startswith("OVA") else "NONE",
    }


lamp3_abundances_df: pd.DataFrame = lamp3_transformed_df

lamp3_features_df: pd.DataFrame = pd.DataFrame.from_records(
    lamp3_transformed_df.index.to_series().apply(determine_lamp3_features),
    index=lamp3_transformed_df.index,
)

output_dir: str = "../lipidome_projector_datasets"

Path.mkdir(Path(output_dir), exist_ok=True)

lamp3_abundances_df.to_csv(output_dir + "/lamp3_abundances.csv")
lamp3_features_df.to_csv(output_dir + "/lamp3_features.csv")
```
