## Supplementary material for "Lipidome visualisation, comparison, and analysis in a vector space": S2 Dataset: yeast_dataset_derivation.py

```
from pathlib import Path

import numpy as np
import pandas as pd

yeast_original_df: pd.DataFrame = pd.read_excel(
    "sd1.xls",
    sheet_name="Lipid species",
    header=None,
)

yeast_transformed_df: pd.DataFrame = (
    yeast_original_df.iloc[8:353, 1:]
    .reset_index(drop=True)
    .set_index(1)
    .rename_axis("Lipid Species", axis="index")
)

yeast_transformed_df = yeast_transformed_df.loc[
    :, yeast_transformed_df.iloc[1] == "Average"
]

yeast_transformed_df.columns = (
    yeast_transformed_df.iloc[0]
    .apply(lambda name: name.replace(" ", "_").replace("°", "").upper()[0:-1])
    .rename("LIPIDOME")
)

yeast_transformed_df.rename(
    columns={"BY4741_24": "WILDTYPE_24", "BY4741_37": "WILDTYPE_37"},
    inplace=True,
)

yeast_transformed_df = (
    yeast_transformed_df.iloc[2:]
    .sort_index(axis="index")
    .sort_index(axis="columns")
)

yeast_transformed_df.replace(0, np.nan, inplace=True)

yeast_transformed_df.dropna(how="all", axis="index", inplace=True)

yeast_transformed_df = yeast_transformed_df.T

dup_col1: pd.Series = yeast_transformed_df.loc[:, "TAG 16:0-16:0-18:0"].iloc[
    :, 0
]
dup_col2: pd.Series = yeast_transformed_df.loc[:, "TAG 16:0-16:0-18:0"].iloc[
    :, 1
]

combined_col: pd.Series = dup_col1.combine_first(dup_col2)

yeast_transformed_df = yeast_transformed_df.drop(columns="TAG 16:0-16:0-18:0")

yeast_transformed_df["TAG 16:0-16:0-18:0"] = combined_col

yeast_abundances_df: pd.DataFrame = yeast_transformed_df

yeast_features_df: pd.DataFrame = (
    yeast_transformed_df.index.to_series()
    .str.split("_", expand=True)
    .rename(
        columns={
            0: "Strain",
            1: "Temperature",
        }
    )
)

output_dir: str = "../lipidome_projector_datasets"

Path.mkdir(Path(output_dir), exist_ok=True)

yeast_abundances_df.to_csv(output_dir + "/yeast_abundances.csv")
yeast_features_df.to_csv(output_dir + "/yeast_features.csv")
```
