## Supplementary material for "Lipidome visualisation, comparison, and analysis in a vector space": S2 Dataset: readme.txt

Supporting Information Dataset 2

- dataset_derivation_scripts:

  This directory contains Python scripts for the extraction of the relevant data
  from the original dataset files and the generation of respective
  "Lipidome Projector" input files. In order to run the scripts to reproduce
  the results, python 3.12 with the packages "pandas", "numpy", "xlrd" and "openpyxl"
  must be installed, and the original datasets must be obtained from the respective
  publications and placed in the directory containing the scripts. The datasets are:

  Drosophila:
  - Publication: Carvalho M, Sampaio JL, Palm W, Brankatschk M, Eaton S, Shevchenko A.
    Effects of diet and development on the Drosophila lipidome. Mol Syst Biol. 2012;8.
    doi:10.1038/msb.2012.29
  - File: msb201229-sup-0002.xls
  - Link to data (June 2024):
    https://www.embopress.org/doi/suppl/10.1038/msb.2012.29/suppl_file/msb201229-sup-0002.xls

  Yeast:
  - Publication: Ejsing CS, Sampaio JL, Surendranath V, Duchoslav E, Ekroos K, Klemm RW, et al.
    Global analysis of the yeast lipidome by quantitative shotgun mass spectrometry.
    Proc Natl Acad Sci U S A. 2009;106. doi:10.1073/pnas.0811700106
  - File: sd1.xls
  - Link to data (June 2024):
    https://www.pnas.org/doi/suppl/10.1073/pnas.0811700106/suppl_file/sd1.xls

  LAMP3:
  - Publication: Lunding LP, Krause D, Stichtenoth G, Stamme C, Lauterbach N, Hegermann J, et al.
    LAMP3 deficiency affects surfactant homeostasis in mice. PLoS Genet. 2021;17.
    doi:10.1371/journal.pgen.1009619
  - File: journal.pgen.1009619.s007.xlsx
  - Link to data (June 2024):
    https://journals.plos.org/plosgenetics/article/file?type=supplementary&id=10.1371/journal.pgen.1009619.s007

- lipidome_projector_datasets:

  This directory contains "Lipidome Projector" input files generated with the scripts
  described above and used for the generation of the lipidome figures (Fig. 4).

- constraints:

  This directory contains fatty acyl ("_fa" suffix) and long-chain base ("_lcb" suffix)
  constraints for the respective "Lipidome Projector" input files used for the generation
  of the lipidome figures (Fig. 4).
