## Supplementary material for "Lipidome visualisation, comparison, and analysis in a vector space": S2 Table

**S2 Table. Stochastic Neighbour Embedding Parameters**

| Optimization | Distance metric | Initialization | Perplexity | Early exaggeration | Iterations |
| --- | --- | --- | --- | --- | --- |
| Barnes-Hut | Cosine | Principal component analysis | 150 | 50 | 1000 |
