## Supplementary material for "Lipidome visualisation, comparison, and analysis in a vector space": S3 Dataset: readme.txt

Supporting Information Dataset 3

This directory contains partially interactive HTML versions of the scatter plots
generated with "Plotly" and used for Fig. 2 (Vector Space (2D)),
S2 Fig. (Vector Space (3D)) and Fig. 4 (Lipidome Dataset Projections).
